# Solanaceae-derived spirostane saponins inhibit dengue virus replication and remodel host-cell metabolism

**DOI:** 10.64898/2026.11.15.766867

**Authors:** Erika Vanessa Jiménez-Posada, Martha Viviana Zuluaga, Oscar Marino Mosquera-Martínez, Diego Durango, Wiston Quiñones, Samuel Vizcaíno-Paez, Sara María Robledo, Marlen Martínez-Gutiérrez

## Abstract

Flaviviruses reprogram host-cell metabolism to support viral replication. In previous work, we showed that Solanaceae-derived spirostane saponins inhibit dengue virus replication and were predicted to interact with viral enzymes. Here, we investigated whether S01, a purified solid from Solanum ovalifolium containing predominantly two characterized spirostane saponins, also modulates host-cell metabolism in association with reduced permissiveness to dengue virus serotype 2 (DENV-2). We performed untargeted GC-QTOF-MS metabolomics in Vero cells under four conditions, including untreated control, S01 treatment, DENV-2 infection, and DENV-2 infection with S01 treatment. Quality-control samples clustered tightly, supporting analytical reproducibility. Principal component analysis revealed separation among experimental groups, and PERMANOVA confirmed significant global metabolic differences (F = 27.567, R² = 0.827, p = 0.001). S01 also retained strong antiviral activity, producing 99.9% inhibition under the metabolomics conditions.

DENV-2 infection was associated with alterations metabolites related to amino acid, carbohydrate, lipid, and sphingolipid metabolism. Pathway analysis highlighted glutamate metabolism, arginine and proline metabolism, glycolysis-related pathways, fatty-acid oxidation and carnitine-associated metabolism, phosphatidylcholine biosynthesis, and sphingolipid metabolism. In uninfected cells, S01 induced a distinct metabolic profile (OPLS-DA R²Y = 0.951, Q² = 0.851; permutation p = 0.002), characterized by increased carbohydrate-related metabolites, including the unresolved Talose/D-glucose feature, gluconic acid, 2-keto-L-gulonic acid, and D-mannitol, together with decreased abundance of selected lipid-related metabolites. During infection, S01 induced a distinct metabolic (OPLS-DA R²Y = 0.928, Q² = 0.841), characterized by lower abundance of several fatty-acid-related and organic-acid metabolites and higher abundance of carbohydrate-related metabolites, while modifying selected DENV-2-associated metabolic alterations.

These findings associate S01 antiviral activity with substantial remodeling of host-cell metabolism. Together with previous docking predictions of interactions with viral targets, the results support a model in which the antiviral activity of Solanaceae-derived spirostane saponins may involve complementary effects on viral and host-associated processes.

## Introduction

Dengue is the most prevalent mosquito-borne viral disease affecting humans, with an estimated 390 million infections annually and nearly half of the global population at risk of exposure (1). Periodic epidemics continue to strain healthcare systems across tropical and subtropical regions, and the clinical spectrum ranges from self-limiting febrile illness to severe manifestations characterized by plasma leakage, coagulopathy, and hypovolemic shock (2). No specific antiviral therapy is currently available, and licensed vaccines provide incomplete protection, with efficacy varying according to factors including viral serotype and prior immune status(3, 4). These limitations underscore the need for new and effective antiviral strategies against DENV infection.

DENV infection is accompanied by extensive remodeling of host-cell metabolism that can support the biosynthetic and energetic demands of viral replication (5). DENV and other flaviviruses have been reported to alter central carbon metabolism and increase glycolytic flux (6, 7). Infection has been associated with increased reliance on fatty acid metabolism (8) and the carnitine shuttle (9), remodeling of phosphatidylcholine and sphingolipid pathways involved in replication-complex assembly (10, 11), and changes in redox homeostasis associated with viral RNA synthesis and virion production (12, 13). These coordinated metabolic changes create a permissive intracellular environment that sustains viral amplification. Consequently, host-directed metabolic modulation has emerged as a complementary antiviral strategy capable of limiting viral replication by restricting access to essential metabolic resources (14).

Natural products represent a valuable source of structurally diverse bioactive compounds with antiviral potential. Steroidal saponins derived from Solanaceae species have demonstrated a range of biological activities, including membrane-interacting (15–17), immunomodulatory (18–20) , and antimicrobial effects (21–23). Their amphipathic structure enables interactions with lipid membranes and may also influence cellular signaling pathways, raising the possibility that they may modulate cellular processes relevant to viral replication. Consistent with this antiviral potential, steroidal saponins isolated from Solanaceae species have been reported to inhibit DENV replication *in vitro* (24) .

In our previous work (25), bioactivity-guided fractionation of *Solanum ovalifolium* led to the isolation of a chromatographically purified solid designated S01, composed predominantly of two structurally characterized spirostane-type steroidal saponins. Compound 1, C1, was identified as 6α-O-[β-D-xylopyranosyl-(1→3)-β-D-quinovopyranosyl]-(25S)-5α-spirostane-3β-ol, whereas compound 2, C2, was identified as neochlorogenin 6-O-α-L-rhamnopyranosyl-(1→3)-β-D-quinovopyranoside. Thus, S01 refers to the purified solid containing predominantly C1 and C2 rather than either saponin individually.

Functional characterization showed that S01 strongly inhibited DENV-2 replication *in vitro* while exerting little detectable effect on viral infectivity following direct virus-compound co-incubation. Notably, pre-treatment of cells with S01 followed by removal of the solid before viral inoculation reduced DENV-2 infection by 99.9%. S01 also retained antiviral activity when administered after viral entry and markedly reduced viral RNA replication. Molecular docking analyses of C1 and C2 further predicted energetically favorable interactions with catalytic regions of DENV NS5 and NS3 (25), suggesting potential interactions with viral proteins, although these predictions do not establish direct binding or enzymatic inhibition. Importantly, the pronounced antiviral activity observed after cellular pre-treatment suggested that interactions with viral proteins may not fully explain the S01-associated antiviral phenotype. Because S01 was removed before viral inoculation under this experimental condition, prior exposure of the may have altered host-cell processes that subsequently influenced DENV infection. This observation raised the possibility that, in addition to potential virus-directed effects, S01 may modulate cellular pathways that contribute to the intracellular environment required for efficient viral replication.

Given the indication that the intracellular context influences S01 efficacy and considering the central role of host metabolism in the DENV replication cycle, we hypothesized that S01 might alter cellular metabolic pathways associated with viral replication. DENV infection requires coordinated remodeling of host-cell metabolism, particularly central carbon, amino acid, and lipid pathways, to support viral biosynthesis, membrane remodeling, and energy production (26, 27). Although the Solanaceae-derived spirostane saponins present in S01 have been predicted to interact with viral enzymes, their effects on host-cell metabolic networks have not been defined.

We therefore investigated whether S01 alters the cellular metabolic state under basal conditions and during DENV-2 infection, and whether these changes are associated with reduced cellular permissiveness to viral replication.

To address this question, we performed untargeted GC-QTOF-MS metabolomics in Vero cells under four experimental conditions: untreated control cells, S01-treated cells, DENV-2-infected cells, and DENV-2-infected cells treated with S01. Multivariate and univariate analyses were integrated with pathway-level analysis to characterize the metabolic profile associated with DENV-2 infection, define S01-associated metabolic changes in uninfected cells, and determine whether S01 modifies infection-associated metabolic remodeling. We found that DENV-2 infection was associated with changes in amino acid, carbohydrate, lipid, and sphingolipid-related metabolism, whereas S01 produced a distinct metabolic profile and modified selected metabolic alterations associated with infection. Collectively, these findings link the antiviral activity of S01 with changes in host-cell metabolism and support a potential host-directed component of its activity, alongside previously proposed interactions with viral targets.

## Materials and Methods

### Steroidal saponins from Solanum ovalifolium

The chromatographically purified solid S01 used in this study was obtained previously from *Solanum ovalifolium* through a bioactivity-guided fractionation and purification workflow. Details of plant collection, voucher information, extraction, fractionation, and chromatographic purification have been described previously (25). Structural characterization of S01 by one- and two-dimensional ^1^H and ^13^C NMR spectroscopy identified two major spirostane-type steroidal saponins. Compound 1, C1, was identified as 6α-O-[β-D-xylopyranosyl-(1→3)-β-D-quinovopyranosyl]-(25S)-5α-spirostane-3β-ol (28, 29), whereas compound 2, C2, was identified as neochlorogenin 6-O-α-L-rhamnopyranosyl- (1→3)-β-D-quinovopyranoside (29). S01 therefore refers to the chromatographically purified solid containing predominantly C1 and C2 and not to either individual compound. The same S01 preparation was used throughout the metabolomic study.

### Cell and virus maintenance

VERO cells (ATCC® CCL-81™), derived from African green monkey (*Cercopithecus aethiops*) kidney epithelium, were maintained in Dulbecco’s modified Eagle medium (DMEM; Gibco®) supplemented with 5% fetal bovine serum (FBS; Gibco®) and 1% antibiotic/antimycotic solution (10 mg/mL streptomycin, 10,000 U/mL penicillin, 0.025 mg/mL amphotericin B) at 37 °C in a 5% CO₂ atmosphere. Antiviral assays were performed using the reference dengue virus strain DENV-2/S16803 (30).

### Antiviral screening *In vitro*

Antiviral and non-cytotoxic concentrations of the enriched solid isolate containing two steroidal saponins were determined using standard cell-based assays as previously described (30, 31). Briefly, Vero cells (3 × 10⁴ cells per well) were seeded in 96-well plates and incubated for 24 h at 37 °C with 5% CO₂ to allow monolayer formation. Cells were treated with the isolated solid at 60 µg/mL for 24 h. Compounds were then removed, and cells were infected with DENV-2/S16803 at a multiplicity of infection (MOI) of 1. After a 2-h adsorption period, the inoculum was removed, cells were washed with 1× PBS, and fresh medium was added. Cultures were incubated for an additional 24 h, after which supernatants were collected for viral quantification by focus-forming assay (FFA), expressed as focus-forming units per milliliter (FFU/mL).

Viral titration was performed by FFA. Vero cells (2 × 10⁴ cells per well) were seeded in 96-well plates and infected with serial dilutions of virus-containing supernatants. At 30 hours post-infection, cells were fixed with 4% paraformaldehyde for 20 min and permeabilized with 0.5% Triton X-100 in PBS for 6 min at room temperature. Immunostaining was carried out using a mouse monoclonal antibody against flavivirus envelope protein (clone 4G2; 1:1000) overnight at 4 °C, followed by incubation with goat anti-mouse IgG–HRP (1:3000; BioLegend, 405306) for 1 h at room temperature. Detection was performed using TrueBlue™ peroxidase substrate (KPL). Infected foci were visualized and counted using a CTL Immunospot analyzer (Cellular Technology Ltd.), and viral titers were calculated as FFU/mL from replicate wells (32).

### Metabolomic experimental design and sample preparation

To evaluate the impact of S01 on host metabolic remodeling during DENV-2 infection, an untargeted intracellular metabolic profiling analysis was conducted using a four-group experimental design: G1 (uninfected, untreated control cells), G2 (cells treated with S01 alone), G3 (cells infected with DENV-2/S16803), and G4 (cells infected and treated with S01). Each condition included six independent biological replicates.

A pre-treatment strategy previously shown to achieve >99.9% inhibition of DENV-2 infection (**Fig. S1**). To obtain sufficient intracellular biomass for metabolite extraction, cell density was increased approximately 30-fold relative to the 96-well antiviral screening format. Vero cells were seeded at 9.0 × 10⁵ cells per well in 6-well plates and incubated for 24 h. Cells assigned to treatment groups were exposed to S01 at 180 µg per well in 1 mL of medium for 24 h. This dose was scaled from the 96-well format to maintain proportional exposure based on an estimated 0.2 ng per cell (30,000 cells treated with 6 µg total in screening format). This calculation was used solely to normalize exposure across experimental formats and should not be interpreted as a measurement of intracellular compound uptake.

Following treatment, infection groups were inoculated with DENV-2/S16803 at a multiplicity of infection (MOI) of 1 for 2 h. Following viral adsorption, the inoculum was removed, fresh medium was added, and cells were incubated for an additional 24 h. Supernatants were collected, and monolayers were rinsed with 500 µL ice-cold PBS. Cells were scraped, pelleted at 250g at 4 °C, using a p, inactivated with 100 µL mass-spectrometry-grade methanol at −20 °C, snap-frozen in liquid nitrogen, and stored at −80 °C. For intracellular metabolite extraction, pellets were resuspended in 500 µL ice-cold 80% methanol, vortexed for 1 min, subjected to three freeze–thaw cycles with 1 min sonication on ice between cycles, and centrifuged at 2264g for 10 min at 4 °C. Supernatants were collected and stored at −80 °C until derivatization.

### GC–QTOF–MS Analysis

For GC–QTOF–MS analysis, 150 µL of extract was dried in a SpeedVac at 35 °C for 1 h. Samples were derivatized with 10 µL methoxyamine hydrochloride in pyridine (15 mg/mL) for 16 h at room temperature in the dark, followed by 10 µL BSTFA containing 1% TMCS for 1 h at 70 °C. After cooling, 60 µL heptane was added prior to injection. Samples were analyzed using an Agilent 7890B gas chromatograph coupled to an Agilent 7250 GC Q-TOF mass spectrometer. One microliter was injected onto an HP-5MS column (30 m × 0.25 mm × 0.25 µm) with helium carrier gas at 0.7 mL/min. The oven program was 60 °C for 1 min, ramped at 10 °C/min to 325 °C, and held for 10 min. Mass spectra were acquired in full-scan mode (m/z 50–600) at 70 eV. Transfer line, ion source, and quadrupole temperatures were 280 °C, 230 °C, and 150 °C, respectively. Raw data were processed using MassHunter Unknowns Analysis B.10.00 for deconvolution and aligned in MassProfiler Professional v15 using a ±0.2 min retention-time window and ±10 ppm mass tolerance. Peak areas were normalized to total ion current. Putative identifications were assigned using the Fiehn GC–MS RTL Library (2011) with retention index matching and a minimum match factor of 60, followed by manual spectral inspection. Only features present in 100% of samples and with <30% coefficient of variation in quality-control replicates were retained (33).

Quality control (QC) samples were generated by pooling equal aliquots of the metabolic extracts from all experimental samples. The pooled QC was initially injected to condition and equilibrate the analytical system and subsequently analyzed after every fourth randomized sample throughout the run. This strategy enabled continuous monitoring of analytical stability, instrument performance, and reproducibility of sample preparation across the entire dataset (34).

## Statistical analysis

Multivariate and univariate analyses were performed in MetaboAnalyst v6.0 (35). Data were mean-centered and scaled prior to analysis. Principal component analysis (PCA) was used to assess global structure and quality-control clustering, and analytical reproducibility based on quality-control (QC) samples. Differences in overall metabolic profiles among experimental groups were assessed using permutational multivariate analysis of variance (PERMANOVA) with 999 permutations.

Pairwise orthogonal partial least squares discriminant analysis (OPLS-DA) models were used to evaluate group separation and identify variables contributing to discrimination between selected experimental groups. Model fit and predictive performance were assessed using R²Y and Q², respectively, together with sevenfold cross-validation and permutation testing (1,000 iterations). Variable importance in projection (VIP) scores ≥1.0 were used to prioritize features contributing to group discrimination rather than as an independent criterion of statistical significance.

Pairwise univariate comparisons were performed using two-sided Student’s t-tests, with P values adjusted for multiple testing using the Benjamini–Hochberg false discovery rate (FDR) method. Statistical significance was defined as an FDR-adjusted P value <0.05. Log₂ fold changes were calculated to describe the direction and magnitude of metabolite differences between groups. Differential abundance was visualized using volcano plots displaying log₂ fold change versus −log₁₀ FDR-adjusted P value (36, 37). Where multivariate and univariate results were integrated, metabolites meeting both VIP ≥1.0 and FDR-adjusted P <0.05 were considered high-priority discriminant features.

Pathway analysis was performed using the the Kyoto Encyclopedia of Genes and Genomes database (KEGG, http://genome.jp/keg) and the Human Metabolome Database (HMDB, https://hmdb.ca/metabolites) to contextualize metabolic alterations associated with DENV-2 infection and S01 treatment.

## Results

We previously demonstrated that solids isolated from *Solanum ovalifolium* exhibit antiviral activity against DENV-2 and other clinically relevant arboviruses (25). Because the antiviral effect was observed when the solid was administered prior to infection, we next investigated whether this effect was associated with measurable changes in the cellular metabolome. We compared the intracellular metabolomic profiles of untreated Vero cells (G1), Vero cells treated with the isolated solid S01 (G2), cells infected with DENV-2/S16803 (G3), and infected cells treated with S01 (G4). Untargeted GC-QTOF-MS metabolomics was performed, and the resulting metabolite profiles were analyzed using multivariate and univariate statistical approaches together with pathway analysis.

We first assessed data quality using an unsupervised principal component analysis (PCA). As shown in **Supplementary Fig. 2**, the quality control (QC) samples formed a compact cluster, supporting analytical reproducibility throughout sample acquisition. The first two principal components explained 62.4% of the total variance (PC1: 44.3%; PC2: 18.1%), and revealed separation among the experimental groups. PERMANOVA analysis confirmed that the separation between groups was statistically significant (F = 27.567; R² = 0.82741; p = 0.001; 999 permutations). Thus, experimental condition accounted for approximately 82.7% of the multivariate variation detected in the dataset. DENV-2-infected cells (G3) showed the greatest displacement along PC1, whereas the infected and S01-treated group (G4) occupied an intermediate position between the G2 and G3 profiles.

### Metabolic alterations induced by DENV-2 infection in Vero cells

To characterize the metabolic perturbations associated with DENV-2 infection, we compared the metabolomic profiles of infected Vero cells (G3) with uninfected controls (G1). Orthogonal partial least-squares discriminant analysis (OPLS-DA) showed separation between the two groups (Fig. 1a). The predicted component accounted 15.7% of the variance in the metabolite matrix, while the model explained 67.2% of the variation associated with class separation (R²Y = 0.672). Cross-validation yielded Q² = 0.315. However, permutation testing (1,000 iterations) yielded p = 0.089, indicating that the supervised model did not reach the conventional threshold for statistical significance. Accordingly, the OPLS-DA separation was interpreted cautiously, and subsequent analyses focused on individual metabolite-level differences.

**Fig. 1.**
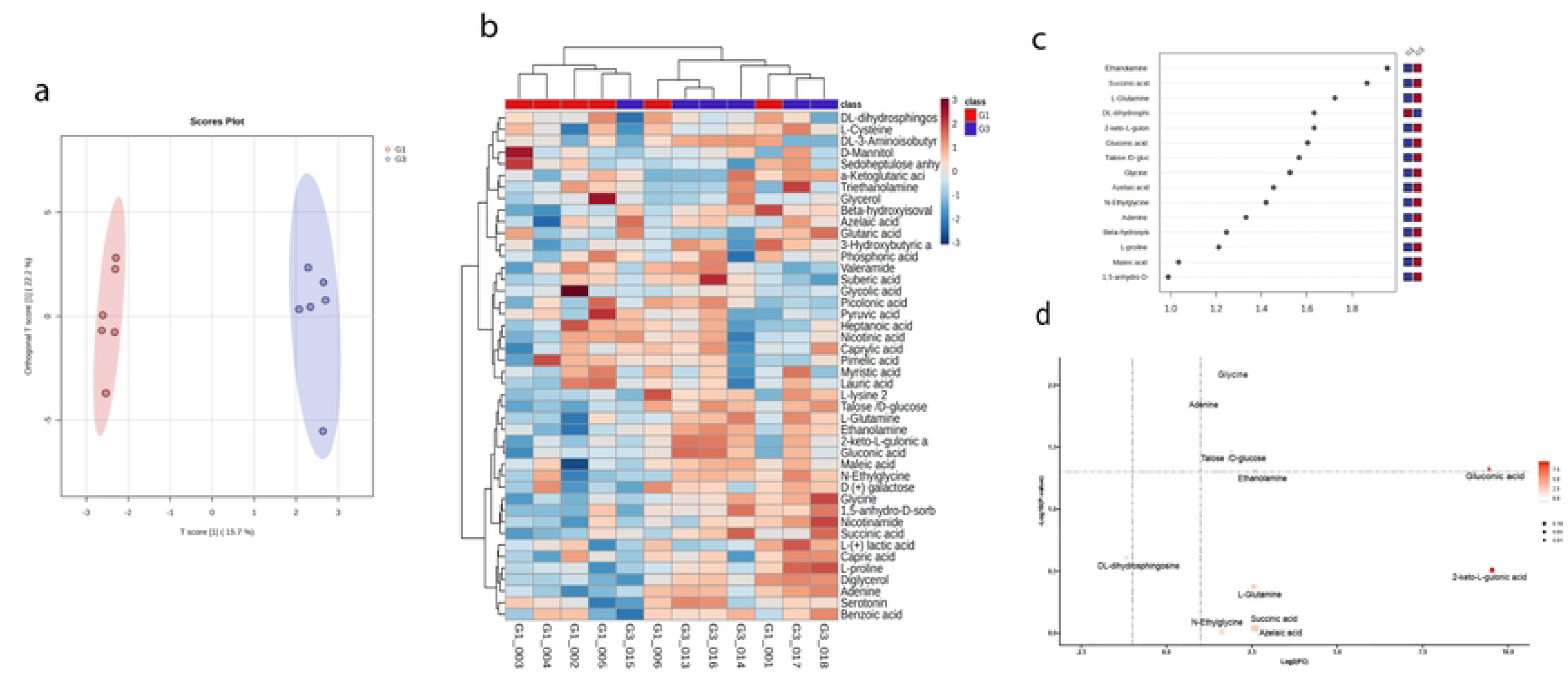
Comparative metabolomic analysis of DENV-2/S16803-infected Vero cells (G3) versus uninfected controls (G1). (a) OPLS-DA score plot showing separation between G1 and G3 (T score [1] = 15.7%; orthogonal T score [1] = 22.2%). (b) Heatmap showing relative abundance patterns of metabolites differing between groups. (c) VIP score plot showing metabolites contributing most strongly to discrimination between G1 and G3. (d) Volcano plot displaying metabolite abundance differences based on log2 fold change and -log10 p-value. All analyses were performed using MetaboAnalyst 6.0.

Univariate analysis identified ten metabolites with nominal abundance differences between G3 and G1 based on unadjusted p < 0.05 (Table 1): ethanolamine, succinic acid, L-glutamine, 2-keto-L-gulonic acid, gluconic acid, DL-dihydrosphingosine (sphinganine), Talose/D-glucose, glycine, azelaic acid, and N-ethylglycine. The heatmap (Fig. 1b) illustrates differences in relative metabolite abundance across biological replicates.

**Table 1.**
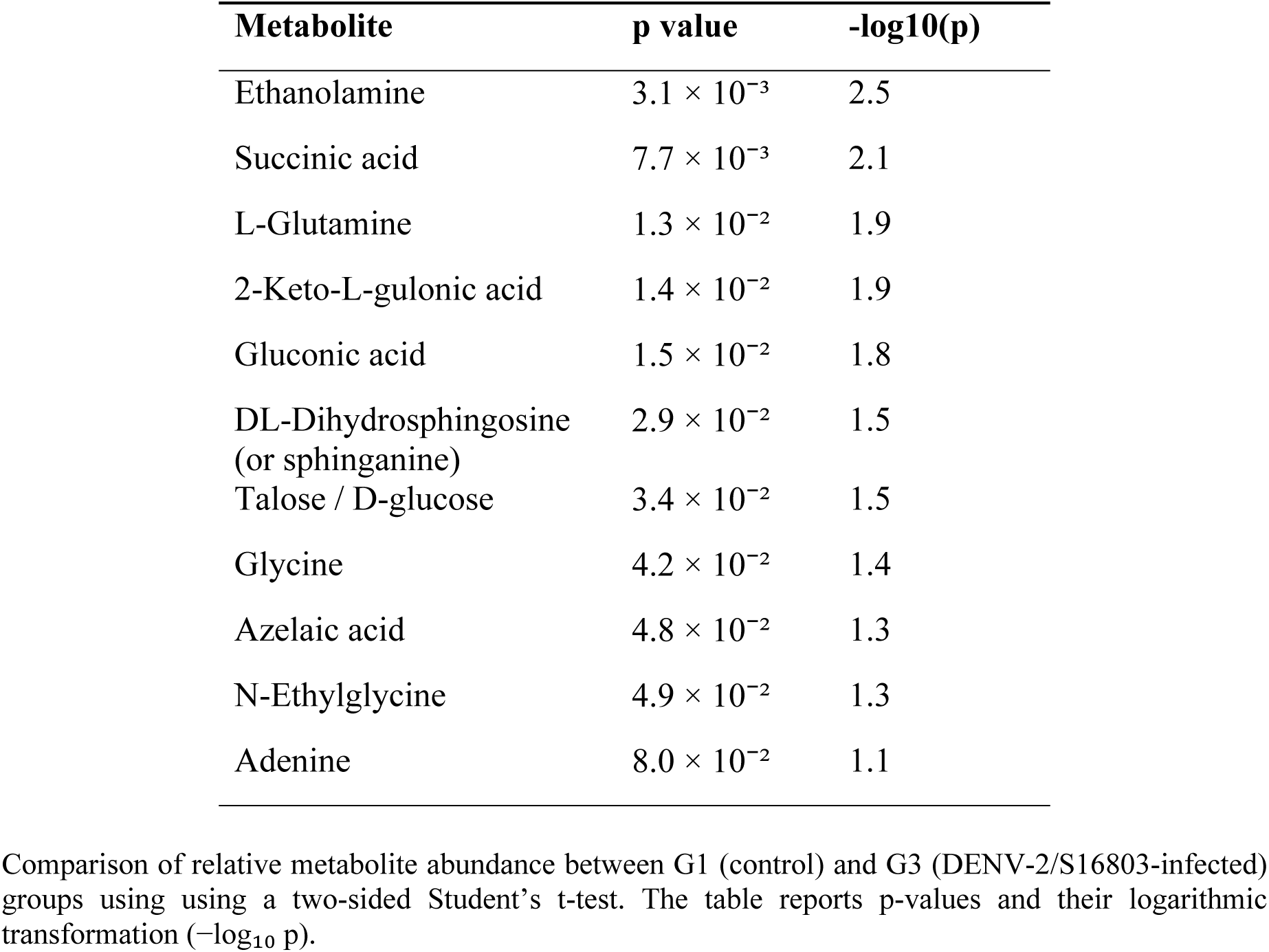
Metabolites reported in the univariate comparison between control cells (G1) and DENV-2/S16803-infected cells (G3)

| **Metabolite** | **p value** | **-log10(p)** |
|---|---|---|
| Ethanolamine | 3.1 × 10<sup>-3</sup> | 2.5 |
| Succinic acid | 7.7 × 10<sup>-3</sup> | 2.1 |
| L-Glutamine | 1.3 × 10<sup>-2</sup> | 1.9 |
| 2-Keto-L-gulonic acid | 1.4 × 10<sup>-2</sup> | 1.9 |
| Gluconic acid | 1.5 × 10<sup>-2</sup> | 1.8 |
| DL-Dihydrosphingosine (or sphinganine) | 2.9 × 10<sup>-2</sup> | 1.5 |
| Talose / D-glucose | 3.4 × 10<sup>-2</sup> | 1.5 |
| Glycine | 4.2 × 10<sup>-2</sup> | 1.4 |
| Azelaic acid | 4.8 × 10<sup>-2</sup> | 1.3 |
| N-Ethylglycine | 4.9 × 10<sup>-2</sup> | 1.3 |
| Adenine | 8.0 × 10<sup>-2</sup> | 1.1 |

Comparison of relative metabolite abundance between G1 (control) and G3 (DENV-2/S16803-infected) groups using using a two-sided Student’s t-test. The table reports p-values and their logarithmic transformation (−log<sub>10</sub> p).

VIP analysis (Fig. 1c) identified metabolites contributing most strongly to the OPLS-DA separation, with the highest scores observed for ethanolamine (VIP 2.01), succinic acid (VIP 1.93), L-glutamine (VIP 1.77), DL-dihydrosphingosine (VIP 1.68), 2-keto-L-gulonic acid (VIP 1.68), gluconic acid (VIP 1.65), and Talose/D-glucose (VIP 1.63). Additional metabolites with VIP > 1 included azelaic acid, N-ethylglycine, adenine, β-hydroxyisovalerate, L-proline, maleic acid, and 1,5-anhydro-D-sorbitol. VIP scores were interpreted as measures of contribution to group discrimination rather than independent evidence of statistical significance.

Pathway analysis using human KEGG annotations (Fig. 2a) identified metabolite sets associated with the DENV-2 infection profile, including glutamate metabolism, arginine and proline metabolism, carnitine-associated metabolism, glycolysis-related pathways, ketone body metabolism, sphingolipid metabolism, phosphatidylcholine biosynthesis, branched-chain fatty-acid oxidation, mitochondrial acetyl-group transfer, and the electron transport chain. A pathway annotated in the database as related to the Warburg effect was also enriched. Overall, these results indicate that DENV-2 infection was associated with measurable changes in metabolites represented across amino acid, carbohydrate, lipid, and sphingolipid-related metabolic pathways.

**Fig. 2.**
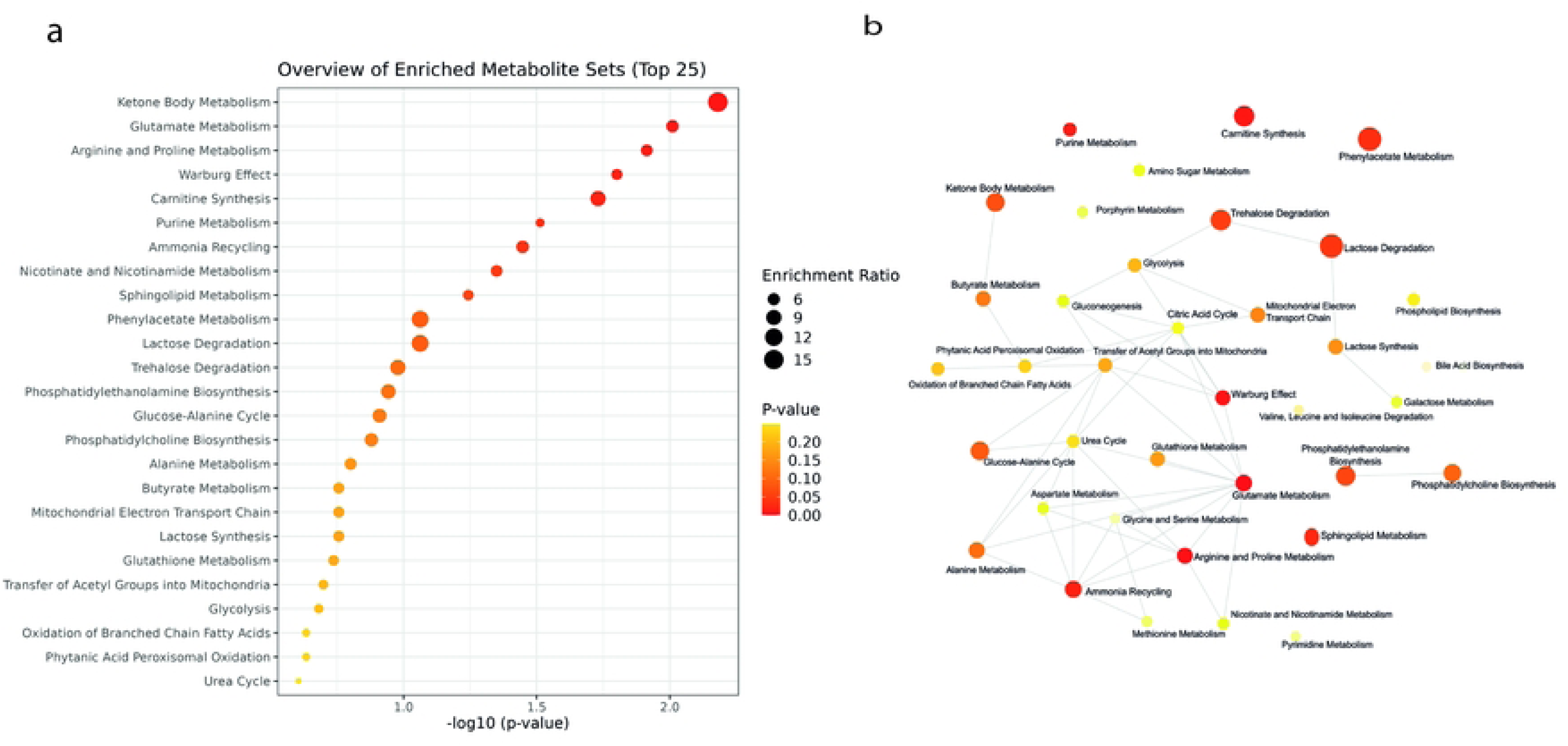
Metabolic pathway analysis associated with DENV-2/S16803 infection (G3 vs. G1). (a) Pathway enrichment plot showing the top 25 enriched pathways ranked by statistical significance (-log₁₀ p value) and enrichment ratio. (b) Metabolic interaction network showing relationships among enriched pathways. All analyses were performed using MetaboAnalyst 6.0.

### Effect of solid S01 on basal cellular metabolism

To determine whether S01 affects cellular metabolism in the absence of viral infection, we compared the metabolomic profiles of untreated control cells (G1) and cells treated with S01 (G2) (**Fig. 3).** OPLS-DA revealed clear separation between the two groups (Fig. 3a). The predictive component accounted for 32.4% of the variation in the metabolomic data, whereas the orthogonal component accounted for an additional 22.4% of non-class-related variation. The model showed R² = 0.951 and Q² = 0.851, and permutation testing (1,000 iterations) yielded p = 0.002, supporting a robust treatment-associated multivariate separation.

**Fig. 3.**
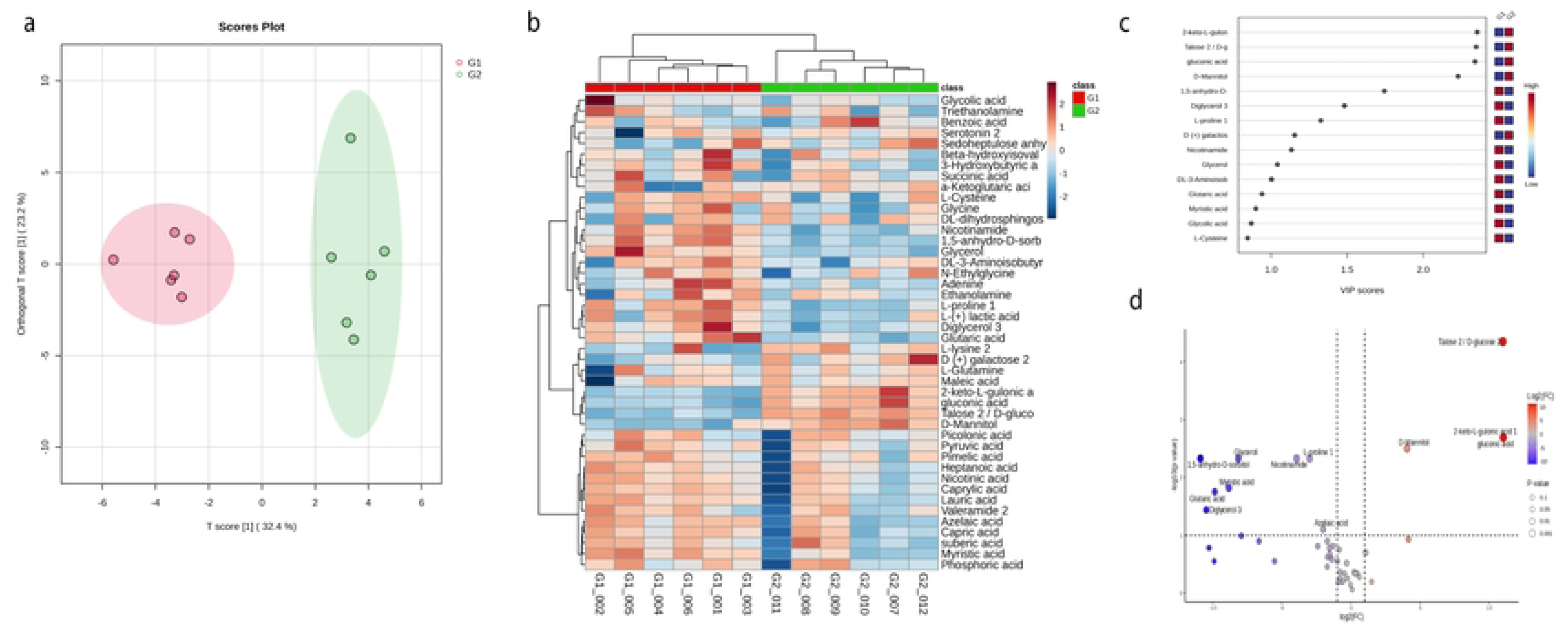
Comparative metabolomic analysis of control cells (G1) and S01-treated cells (G2). (a) OPLS-DA score plot showing separation between G1 and G2 (R² = 0.951; Q² = 0.851; permutation test p = 0.002). The predictive and orthogonal components accounted for 32.4% and 22.4% of the modeled variation, respectively. (b) Heatmap showing relative abundance patterns between groups. (c) VIP score plot showing metabolites contributing most strongly to group discrimination. (d) Volcano plot showing metabolite abundance differences according to log2 fold change and -log10 p-value (p < 0.05). All analyses were performed using MetaboAnalyst 6.0.

The heatmap (Fig. 3b) showed distinct relative-abundance patterns between G1 and G2. S01-treated cells showed higher abundance of several carbohydrate-related metabolites, including Talose/D-glucose, gluconic acid, 2-keto-L-gulonic acid, and D-mannitol. In contrast, nicotinamide, 1,5-anhydro-D-sorbitol, glycerol, glutaric acid, L-proline, L-lactic acid, and diglycerol were more abundant in untreated cells.

VIP analysis (Fig. 3c) identified Talose/D-glucose, gluconic acid, 2-keto-L-gulonic acid, and D-mannitol among the strongest contributors to discrimination between G1 and G2 (VIP > 1.6), followed by L-proline, nicotinamide, 1,5-anhydro-D-sorbitol, and glutaric acid. Univariate analysis (Table 2) identified pronounced abundance differences in several carbohydrate-related metabolites, including Talose/D-glucose (p = 9.92 × 10⁻⁷), gluconic acid (p = 9.54 × 10⁻⁵), 2-keto-L-gulonic acid (p = 1.4 × 10⁻⁴), and D-mannitol (p = 2.8 × 10⁻⁴). Conversely, several metabolites, including 1,5-anhydro-D-sorbitol, glycerol, myristic acid, and glutaric acid, showed lower abundance following S01 exposure. Together, the multivariate and univariate analyses indicate that S01 treatment is associated with a distinct basal metabolic profile characterized by increased abundance of selected carbohydrate-related metabolites and decreased abundance of several lipid-, organic-acid-, and amino-acid-related metabolites.

**Table 2.**
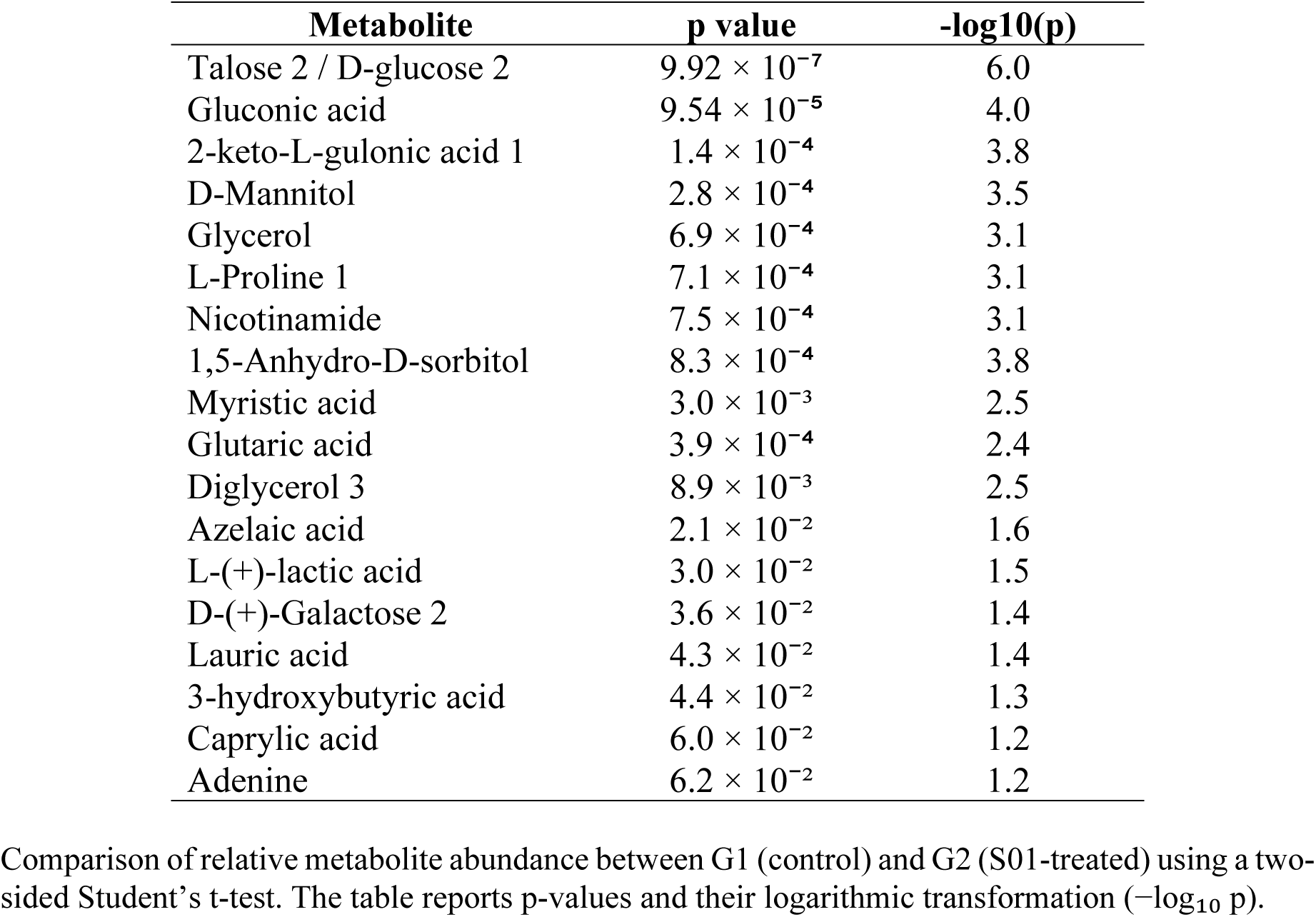
Differentially expressed metabolites between control cells (G1) and S01-treated cells (G2)

| Metabolite | p value | -log10(p) |
|---|---|---|
| Talose 2 / D-glucose 2 | 9.92 × 10<sup>−7</sup> | 6.0 |
| Gluconic acid | 9.54 × 10<sup>−5</sup> | 4.0 |
| 2-keto-L-gulonic acid 1 | 1.4 × 10<sup>−4</sup> | 3.8 |
| D-Mannitol | 2.8 × 10<sup>−4</sup> | 3.5 |
| Glycerol | 6.9 × 10<sup>−4</sup> | 3.1 |
| L-Proline 1 | 7.1 × 10<sup>−4</sup> | 3.1 |
| Nicotinamide | 7.5 × 10<sup>−4</sup> | 3.1 |
| 1,5-Anhydro-D-sorbitol | 8.3 × 10<sup>−4</sup> | 3.8 |
| Myristic acid | 3.0 × 10<sup>−3</sup> | 2.5 |
| Glutaric acid | 3.9 × 10<sup>−4</sup> | 2.4 |
| Diglycerol 3 | 8.9 × 10<sup>−3</sup> | 2.5 |
| Azelaic acid | 2.1 × 10<sup>−2</sup> | 1.6 |
| L-(+)-lactic acid | 3.0 × 10<sup>−2</sup> | 1.5 |
| D-(+)-Galactose 2 | 3.6 × 10<sup>−2</sup> | 1.4 |
| Lauric acid | 4.3 × 10<sup>−2</sup> | 1.4 |
| 3-hydroxybutyric acid | 4.4 × 10<sup>−2</sup> | 1.3 |
| Caprylic acid | 6.0 × 10<sup>−2</sup> | 1.2 |
| Adenine | 6.2 × 10<sup>−2</sup> | 1.2 |

Comparison of relative metabolite abundance between G1 (control) and G2 (S01-treated) using a two-sided Student’s t-test. The table reports p-values and their logarithmic transformation (−log<sub>10</sub> p).

Pathway analysis of metabolites associated with S01 treatment (Fig. 4a) highlighted pathways related to carbohydrate metabolism, including lactose degradation, galactose metabolism, glycolysis, and gluconeogenesis-related processes. Lipid-associated pathways included fatty-acid biosynthesis, sphingolipid metabolism, and very-long-chain fatty-acid β-oxidation, whereas amino-acid-related pathway annotations included carnitine synthesis and lysine, arginine, and proline metabolism. The analysis also identified a database-defined pathway associated with the Warburg effect. These results indicate that S01 exposure is associated with broad changes in metabolites represented across carbohydrate, lipid, and amino acid metabolic pathways.

**Fig. 4.**
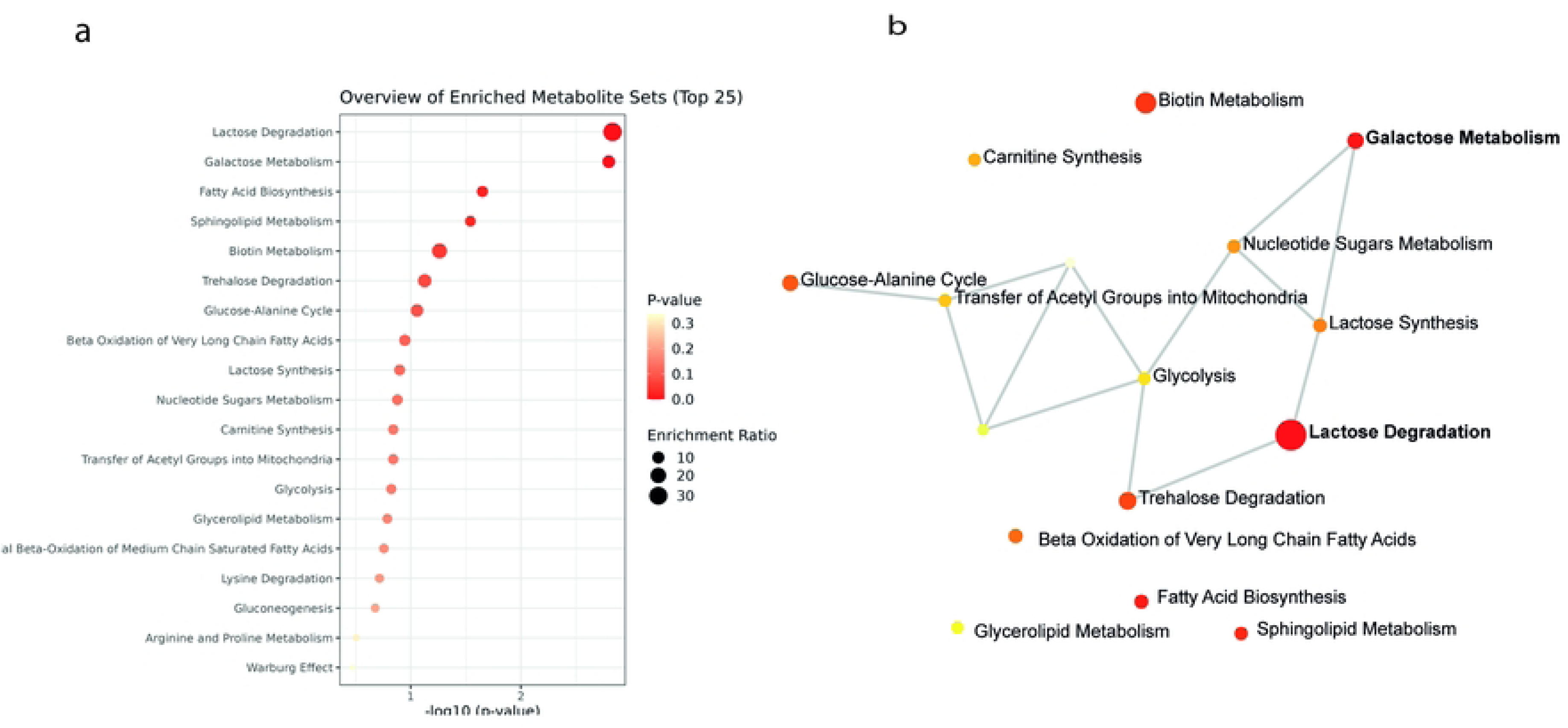
Metabolic pathway analysis of S01-induced metabolic reprogramming (G2 vs. G1). (a) Pathway analysis plot showing the top 25 enriched metabolic pathways. ranked by statistical significance (−log₁₀ p-value) and enrichment ratio. (b) Metabolic interaction network showing associations among enriched pathway annotations. All analyses were performed using MetaboAnalyst 6.0.

### Effect of S01 antiviral treatment on cellular metabolism during DENV-2 infection

We next examined the effect of solid S01 on infected Vero cells. Due to the limited amount of material available, the previously described antiviral assay was scaled to a higher cell-density format, as detailed in the methods section. The dose adjustment was validated experimentally, since comparable antiviral activity was observed in both formats, with 99.9% inhibition for S01 and 91.4% inhibition for Celgosivir (p < 0.001) in the 96-well and 6-well systems, respectively (**Supplementary Fig. 1**).

The multivariate OPLS-DA analysis revealed distinct intracellular metabolomic profiles between DENV-2/S16803-infected cells (G3) and infected cells treated with S01 (G4). The model generated one predictive component that explained 35.4% of the variance in the metabolite matrix and 92.8% of the variance associated with group separation (R²Y). The model showed high predictive performance with Q² = 0.841, while the orthogonal component accounted for an additional 27.1% of non-class-related variation. Permutation testing with 1,000 iterations yielded p = 0.002, supporting the statistical significance of the separation between infection alone and infection with S01 treatment (**Fig. 5a**).

**Fig. 5.**
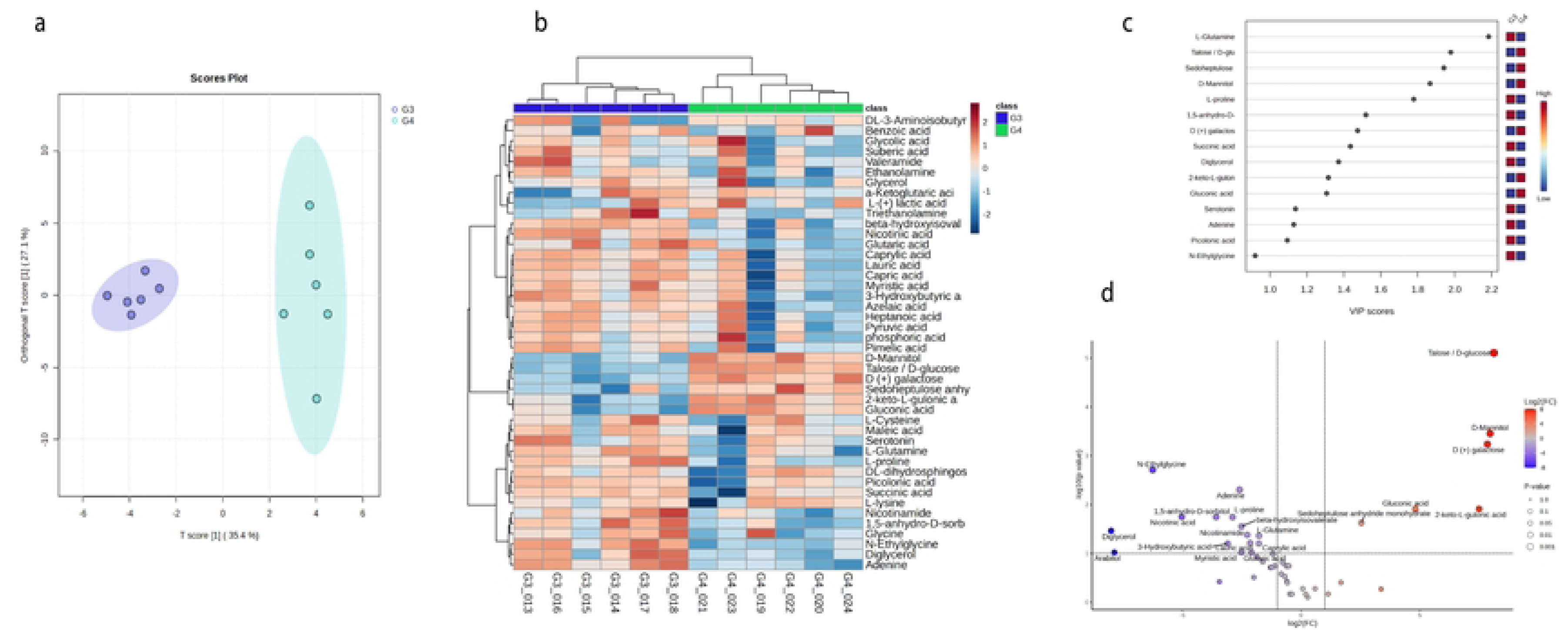
Comparative metabolomic analysis between DENV-2/S16803-infected cells and infected cells treated with S01. **a**, OPLS-DA score plot showing separation between G3 and G4, with a predictive component explaining 35.4% of the metabolite variance and an orthogonal component explaining 27.1% of non-class-related variation. The model showed R²Y = 0.928 and Q² = 0.841, with permutation testing yielding p = 0.002. **b**, Heatmap showing relative abundance patterns of metabolites differing between G3 and G4. **c**, VIP score plot from the OPLS-DA model showing metabolites contributing most strongly to group discrimination. **d**, Volcano plot showing differential metabolite abundance between G3 and G4 based on log₂ fold change and −log₁₀ FDR-adjusted p-value. Positive log₂ fold-change values indicate higher relative abundance in G4, whereas negative values indicate lower abundance relative to G3. Dashed vertical lines indicate |log₂FC| > 1, and the horizontal line indicates FDR < 0.05.

The heatmap (Fig. 5b) showed distinct relative-abundance patterns between G3 and G4. Talose/D-glucose, D-mannitol, D-galactose, 2-keto-L-gulonic acid, and gluconic acid were more abundant in S01-treated infected cells. In contrast, diglycerol, glutaric acid, myristic acid, lauric acid, caprylic acid, and nicotinamide were more abundant in infected cells without S01 treatment. Thus, S01 treatment was associated with bidirectional changes in metabolite abundance rather than a uniform increase or decrease across the metabolome.

Relative to G3, S01-treated infected cells showed lower abundance of several lipid- and organic-acid-related metabolites, including myristic, lauric, and caprylic acids, glutaric acid, and diglycerol. Conversely, S01 treatment was associated with higher abundance of several carbohydrate-related metabolites, sugar acids, and polyols, including Talose/D-glucose, D-galactose, D-mannitol, the sedoheptulose-related feature, 2-keto-L-gulonic acid, and gluconic acid.

The FDR-adjusted volcano plot (Fig. 5d) further illustrated the magnitude and direction of the differences between G3 and G4. Talose/D-glucose showed the largest positive change (log2FC = 8.13; adjusted p = 7.87 × 10⁻⁶), followed by D-mannitol (log2FC = 7.96; adjusted p = 3.52 × 10⁻⁴), D-galactose (log2FC = 7.85; adjusted p = 5.83 × 10⁻⁴), 2-keto-L-gulonic acid (log2FC = 7.50; adjusted p = 0.0123), and gluconic acid (log2FC = 4.81; adjusted p = 0.0123). In contrast, N-ethylglycine, adenine, nicotinic acid, 1,5-anhydro-D-sorbitol, L-proline, β-hydroxyisovalerate, diglycerol, nicotinamide, and L-glutamine showed negative log2 fold changes, indicating lower relative abundance in G4.

Univariate analysis using two-sided Student’s t-tests identified 15 metabolites nominal abundance differences between G3 and G4 based on unadjusted p < 0.05 (Table 3). The strongest differences were observed for Talose/D-glucose (p = 1.75 × 10⁻⁷), D-mannitol (p = 1.57 × 10⁻⁵), D-galactose (p = 3.89 × 10⁻⁵), N-ethylglycine (p = 1.73 × 10⁻⁴), and adenine (p = 5.54 × 10⁻⁴). FDR-adjusted p values for metabolites that were additionally represented in the heatmap and contributed to group discrimination with VIP > 1 are reported in Supplementary Table 1. Together, the univariate and multivariate analyses support a pronounced S01-associated metabolic shift during DENV-2 infection.

**Table 3.** Differentially expressed metabolites between DENV-2/S16803-infected cells (G3) and infected cells treated with S01 (G4).

| **Metabolite** | **p value** | **-log10(p)** |
|---|---|---|
| Talose / D-glucose | 1.7 × 10<sup>−7</sup> | 6.8 |
| D-Mannitol | 1.6 × 10<sup>−5</sup> | 4.8 |
| D-(+)-Galactose | 3.9 × 10<sup>−5</sup> | 4.4 |
| N-Ethylglycine | 1.7 × 10<sup>−4</sup> | 3.8 |
| Adenine | 5.5 × 10<sup>−4</sup> | 3.3 |
| 2-keto-L-gulonic acid | 1.8 × 10<sup>−3</sup> | 2.7 |
| Gluconic acid | 1.9 × 10<sup>−3</sup> | 2.7 |
| Nicotinic acid | 3.2 × 10<sup>−3</sup> | 2.5 |
| 1,5-Anhydro-D-sorbitol | 3.6 × 10<sup>−3</sup> | 2.4 |
| L-Proline | 4.0 × 10<sup>−3</sup> | 2.4 |
| Sedoheptulose anhydride monohydrate | 5.8 × 10<sup>−3</sup> | 2.2 |
| β-Hydroxyisovalerate | 8.0 × 10<sup>−3</sup> | 2.1 |
| Diglycerol | 1.0 × 10<sup>−2</sup> | 2.0 |
| Nicotinamide | 1.3 × 10<sup>−2</sup> | 1.9 |
| L-Glutamine | 1.5 × 10<sup>−2</sup> | 1.8 |

Pathway enrichment analysis (**Fig. 6a**) identified several metabolite sets associated with S01 treatment during DENV-2 infection. These included carnitine-associated metabolism and fatty acid β-oxidation, glycolysis, phosphatidylcholine biosynthesis, ketone body metabolism, sphingolipid metabolism, and pathways involving arginine and proline metabolism. These pathways encompass processes related to cellular bioenergetics, lipid metabolism, membrane-associated metabolism, and redox-related processes. Higher −log₁₀(p) values reflected stronger statistical enrichment, while the enrichment ratio represented the relative representation of metabolites from each pathway in the dataset. These results provide metabolic context for the observed metabolite differences but do not by themselves establish changes in pathway flux or activity.

**Fig. 6.**
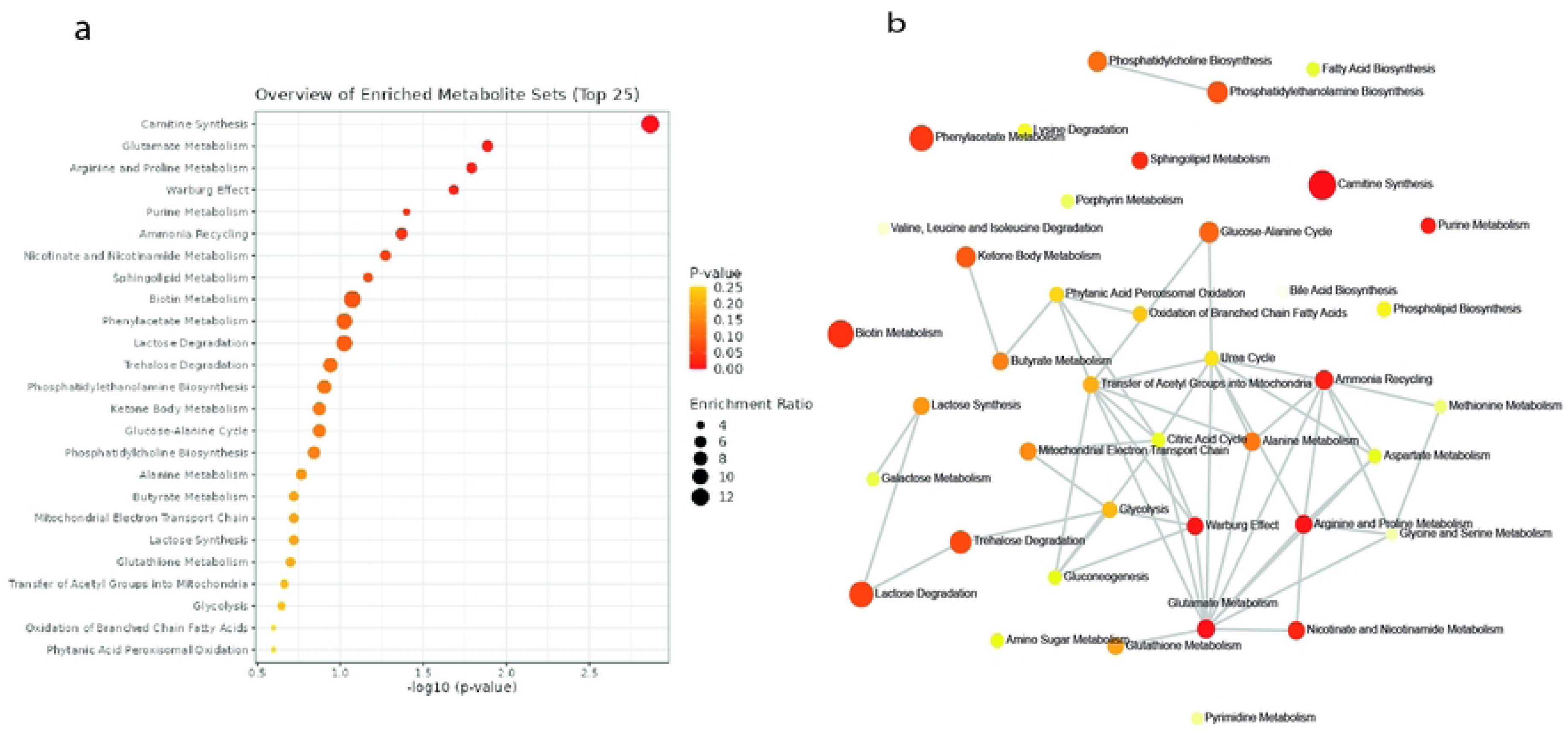
Metabolic pathway analysis of S01 antiviral treatment during DENV-2/S16803 infection (G4 vs. G3). (a) Pathway analysis plot showing the top 25 enriched metabolite sets, ranked by statistical significance (-log10 p-value) and enrichment ratio. (b) Metabolic interaction network showing associations among enriched pathway annotations. All analyses were performed using MetaboAnalyst 6.0.

Collectively, these data indicate that DENV-2 infection induces a measurable metabolic reprogramming in Vero cells, characterized by coordinated alterations in amino acid metabolism, carbohydrate utilization, and sphingolipid-associated pathways. In uninfected cells, S01 establishes a distinct metabolic state marked by increased acidic sugars and polyols linked to redox-associated pathways and reduced lipid intermediates.

Importantly, during DENV-2 infection, S01 shifted the intracellular metabolome away from the infection-associated metabolic phenotype. This shift was characterized by the partial reversal of selected DENV-2-associated metabolic alterations, accompanied by additional treatment-specific changes affecting carbohydrate-, lipid-, and amino acid-related metabolites. These findings suggest that the antiviral activity of S01 is associated with extensive remodeling of host-cell metabolism and support a potential role for host-directed metabolic modulation in its mechanism of action.

## Supplementary figure legends

Supplementary Fig. 1. Inhibition of DENV-2/S16803 infection under the experimental conditions used for metabolomic analysis. Infection percentages in S01-treated cultures (0.2 ng/cell; 60 µg/mL) were determined by quantification of infectious viral particles in supernatants using a focus-forming assay. Untreated cells (medium only) were considered 100% infection, and celgosivir (77 µM) was included as an inhibition control. Asterisks indicate statistically significant differences compared with untreated controls (*p < 0.05; **p < 0.01; ***p < 0.001; one-way ANOVA with Dunnett’s post hoc test; n = 6).

Supplementary Fig. 2. Principal component analysis (PCA) of all experimental and QC samples. The plot shows the distribution of G1, G2, G3, and G4 together with QC samples. The first two principal components explain 62.4% of the total variance (PC1, 44.3%; PC2, 18.1%). PERMANOVA confirmed significant differences among groups (F = 27.567; R² = 0.82741; p = 0.001; 999 permutations), indicating that experimental group accounted for 82.74% of the multivariate variation in the dataset. Tight clustering of QC samples supports analytical reproducibility.

## Discussion

In this study, we investigated the antiviral activity of the purified solid S01 and its effects on the intracellular metabolome of Vero cells infected with DENV-2/S16803, building directly on our previous bioprospecting work on Solanaceae species from Colombia’s Eje Cafetero region aimed at identifying antiviral candidates against DENV, ZIKV, and CHIKV (25). In that study, bioactivity-guided isolation from *Solanum ovalifolium* led to the identification and structural characterization of S01 as a spirostane-saponin-containing solid with potent antiviral activity against DENV-2. Molecular docking of its major constituents further predicted interactions with catalytic regions of DENV NS3 and NS5. Building on these findings, the present study addressed an unresolved question: whether the antiviral phenotype of S01 is also associated with alterations in host-cell metabolism during DENV-2 infection.

To investigate the cellular context associated with this antiviral phenotype, we performed untargeted GC-QTOF-MS metabolomic profiling under four experimental conditions: uninfected control cells (G1), S01-treated cells (G2), DENV-2-infected cells (G3), and infected cells treated with S01 (G4). This experimental design allowed us to distinguish metabolic changes associated with viral infection, S01 exposure in uninfected cells, and S01 treatment during infection.

DENV-2 infection was associated with alterations in host-cell metabolism involving amino acid-, carbohydrate-, lipid-, and sphingolipid-related metabolites. Infected cells showed altered abundance of TCA-cycle intermediates, including succinate and α-ketoglutarate, as well as amino acid-derived metabolites and sphingolipid precursors. Changes in medium-chain fatty acids and dihydrosphingosine were also observed, consistent with perturbations in lipid-associated metabolism and with the established dependence of flaviviruses on host membrane remodeling during replication-complex formation (7, 26, 36). However, because the G1-versus-G3 OPLS-DA model showed only moderate predictive performance and did not reach significance by permutation testing, these infection-associated metabolic patterns should be interpreted cautiously and in conjunction with the corresponding univariate and pathway-level analyses.

In the absence of viral infection, S01 was associated with a distinct metabolic phenotype characterized by increased abundance of several carbohydrate-related metabolites and polyols, including Talose/D-glucose, gluconic acid, 2-keto-L-gulonic acid, and D-mannitol, together with decreased abundance of selected lipid- and amino acid-related metabolites. The increase in carbohydrate-related metabolites and oxidized sugar derivatives suggests remodeling of carbohydrate-associated metabolism under S01 treatment. Carbohydrate metabolic pathways are closely integrated with cellular redox homeostasis, particularly through NADPH-generating systems such as the pentose phosphate pathway. Thus, the observed profile is consistent with altered carbohydrate- and redox-associated metabolism, although the specific contribution of these metabolites to cellular antioxidant homeostasis remains to be established.

The concurrent decrease in selected lipid-related metabolites is also consistent with remodeling of lipid-associated metabolism. Given the dependence of flaviviruses on host lipid and bioenergetic resources, these changes raise the possibility that S01 exposure may alter the cellular metabolic environment in a manner that is less permissive to viral replication (37). However, the present steady-state metabolomics data do not establish whether S01 directly alters lipid biosynthesis, β-oxidation, metabolic flux, or substrate availability for viral replication. Targeted metabolomics and isotope-tracing studies will therefore be required to determine whether the observed changes reflect functional alterations in lipid utilization or other metabolic processes relevant to DENV replication

During DENV-2 infection, S01 further modified the intracellular metabolome. Relative to infected cells without treatment, S01-treated infected cells showed lower abundance of several fatty acids and dicarboxylic acids, together with higher abundance of carbohydrate-related metabolites, including Talose/D-glucose, 2-keto-L-gulonic acid, gluconic acid, D-mannitol, and D-galactose. Importantly, S01 treatment did not simply restore the metabolome to the uninfected state; rather, it modified selected infection-associated alterations while also producing treatment-specific metabolic changes. These metabolic differences occurred under S01 treatment conditions previously shown to be non-cytotoxic, supporting their interpretation as treatment-associated alterations rather than nonspecific consequences of overt cell damage (38).

The lipid-associated changes observed during S01 treatment are compatible with altered host lipid metabolism and may be relevant to processes involved in flavivirus replication. Given the established dependence of flavivirus replication on host membrane remodeling, these changes may be relevant to replication-complex formation (39); however, they do not by themselves demonstrate inhibition of this process. Likewise, the observed metabolic changes occurred alongside profound suppression of viral replication the increased abundance of several carbohydrate-related metabolites is compatible with altered carbon utilization and redox-associated metabolism but does not establish increased pathway flux or a specific redox mechanism. Nevertheless, the concurrence of S01-associated metabolic remodeling with strong inhibition of viral replication supports the hypothesis that host metabolic effects may contribute to its antiviral activity.

A central consideration in interpreting these findings is that suppression of viral replication can itself alter the host-cell metabolome. Consequently, the present experimental design cannot fully distinguish primary metabolic effects directly induced by S01 from secondary metabolic consequences of reduced viral burden. The observed metabolic phenotype should therefore be interpreted as being associated with S01-mediated antiviral activity rather than as evidence that metabolic remodeling is itself the causal mechanism of viral inhibition.

Several additional limitations warrant consideration. The experiments were conducted in Vero cells, which lack a fully functional interferon response. Validation in human primary cells and other immunocompetent models will therefore be necessary to determine the physiological relevance and generalizability of the observed metabolic responses. In addition, the present untargeted metabolomics approach measures relative steady-state metabolite abundance and cannot establish metabolic flux or identify the specific host targets responsible for the S01-associated phenotype. Future studies integrating targeted metabolomics, lipidomics, and stable-isotope tracing, including ¹³C-based approaches, will be important for determining whether S01 directly alters specific metabolic reactions, enzymatic activities, substrate utilization, or upstream regulatory pathways (40).

In conclusion, this study shows that the antiviral activity of the *Solanum ovalifolium*-derived solid S01 is accompanied by substantial changes in host-cell metabolism. DENV-2 infection was associated with alterations in amino acid-, carbohydrate-, lipid-, and sphingolipid-related metabolism, whereas S01 produced a distinct metabolic profile in uninfected cells and substantially modified the intracellular metabolome during infection. Selected DENV-2-associated metabolic alterations were modified in S01-treated infected cells, while additional treatment-associated changes occurred in parallel with strong antiviral activity. Together with previous molecular docking predictions involving DENV NS3 and NS5, these findings support the hypothesis that the antiviral phenotype of S01 may involve complementary virus- and host-associated effects. However, the present data do not establish a causal role for metabolic remodeling in antiviral activity. Validation in primary human cells, together with targeted metabolomics and flux-based studies, will be required to determine the mechanistic contribution of host metabolic changes to S01-mediated inhibition of DENV-2.

## Acknowledgements

We sincerely thank the Vice-Rectorate for Research, Innovation and Extension at Universidad Tecnológica de Pereira (UTP) for its administrative and institutional support throughout the development of this project. We also thank the GB-PN group at Universidad Tecnológica de Pereira and the QOPN group at Universidad de Antioquia for providing access to facilities and for their scientific and technical support. We are particularly grateful to Dr. Eva Harris, Division of Infectious Diseases and Vaccinology, School of Public Health, University of California, Berkeley, for providing reagents, logistical and software resources, as well as for her scientific guidance and continued support. We also sincerely thank Dr. Monica Cala and the MetCore team from Universidad de los Andes for their support with the metabolomics analyses, which was essential for the development of this component of the study.

